# Heterozygous *RFX6* protein truncating variants are associated with Maturity-Onset Diabetes of the Young (MODY) with reduced penetrance

**DOI:** 10.1101/101881

**Authors:** Kashyap A Patel, Jarno Kettunen, Markku Laakso, Alena Stančáková, Thomas W Laver, Kevin Colclough, Matthew B. Johnson, Marc Abramowicz, Leif Groop, Päivi J. Miettinen, Maggie H Shepherd, Sarah E Flanagan, Sian Ellard, Nobuya Inagaki, Andrew T Hattersley, Tiinamaija Tuomi, Miriam Cnop, Michael N Weedon

## Abstract

Finding new genetic causes of monogenic diabetes can help to understand development and function of the human pancreas. We aimed to find novel protein–truncating variants causing Maturity–Onset Diabetes of the Young (MODY), a subtype of monogenic diabetes. We used a combination of next–generation sequencing of MODY cases with unknown aetiology along with comparisons to the ExAC database to identify new MODY genes. In the discovery cohort of 36 European patients, we identified two probands with novel *RFX6* heterozygous nonsense variants. *RFX6* protein truncating variants were enriched in the MODY discovery cohort compared to the European control population within ExAC (odds ratio, OR=131, P=l×l0^‐4^). We found similar results in non–Finnish European (n=348, OR=43, P=5×l0^‐5^) and Finnish (n=80, OR=22, P=1×l0^‐6^) replication cohorts. The overall meta–analysis OR was 34 (P=l×l0^‐16^). *RFX6* heterozygotes had reduced penetrance of diabetes compared to common *HNF1A* and *HNF4A*–MODY mutations (27%, 70% and 55% at 25 years of age, respectively). The hyperglycaemia resulted from beta–cell dysfunction and was associated with lower fasting and stimulated gastric inhibitory polypeptide (GIP) levels. Our study demonstrates that heterozygous *RFX6* protein truncating variants are associated with MODY with reduced penetrance.

---

Finding the genetic cause of rare familial diabetes (monogenic diabetes) provides new biological insights into human pancreas development and function, as well as potentially novel therapeutic targets with important treatment implications^1^. Maturity–Onset Diabetes of the Young (MODY) is monogenic diabetes resulting from beta–cell dysfunction which usually present before the age of 25 years in non–obese patients who are non–insulin dependent and have an autosomal dominant inheritance of diabetes^2^. Mutations in *HNF1A, HNF4A* and *GCK* are the commonest causes of MODY responsible for ~60 % of MODY aetiology^1^.

There has been limited recent success in finding new MODY genes. *WFS1* heterozygous variants and loss–of–function variants in *theAPPL1* gene were shown to be a rare cause of MODY^3,4^. The reason for this limited success is the difficulty of distinguishing monogenic diabetes patients from those with type 1 diabetes^5,6^, or from the increasing number of patients with early–onset type 2 diabetes due to rising rates of obesity. Another important reason is the lack of large pedigrees with an autosomal dominant pattern of inheritance of diabetes which would allow classical linkage analysis to be performed and which was used to discover the most common forms of MODY such as *GCK, HNF1A* and *HNF4A*^7-10^.

Rare–variant association testing is an important step to confirm the pathogenicity of novel variants in monogenic disease^11^. Rare–variant association testing particularly for comparing the frequency of novel protein–truncating variants (PTVs) in monogenic cases with unknown aetiology to the frequency in large control cohorts is now possible because of the availability of resources such as ExAC–a database of protein coding variants in large control populations^12^. This allows burden testing of the frequency of novel or rare coding variants in diseases of interest and a comparison to rates in controls to identify new genetic causes of monogenic disease.

In this study, we undertook next–generation sequencing of MODY cases with unknown aetiology and compared the frequency of PTVs to large publicly available control cohorts to identify new MODY genes. Our study showed that heterozygous *RFX6* PTVs are associated with MODY.

## Results

### Heterozygous *RFX6* PTVs were identified in MODY patients with unknown aetiology

To identify patients with novel heterozygous PTVs, we first assessed 38 European (non–Finnish) probands with a strong MODY–like phenotype who did not have mutations in the common MODY genes *(GCK, HNF1A, HNF4A)* by Sanger sequencing (**Supplementary Table 1**).To exclude the other known/less common causes of monogenic diabetes, these patients underwent comprehensive targeted–next generation sequencing (NGS) for all 29 known monogenic diabetes genes, including genes for neonatal diabetes, MODY and mitochondrial diabetes, lipodystrophy or other forms of syndromic diabetes^13^ (**Supplementary Table 2**).We identified two probands with mutations in the known MODY gene *HNF1B*^13,14^. The analysis of heterozygous PTVs in the 29 genes on the targeted panel identified two unrelated probands with a novel heterozygous nonsense variant in *Regulatory Factor X 6 [RFX6*) (Family 1–p.Leu292Ter, Family 2–p.Lys351Ter) (Table 1, Fig. 1, **Supplementary Table 3**). We did not identify any rare (<1%) missense *RFX6* variants in this cohort. *RFX6* was part of the targeted sequencing panel because recessive *RFX6* variants (missense and/or protein–truncating) are a known cause of syndromic neonatal diabetes^15^, but heterozygotes were not previously known to have any phenotype.

**Figure 1.**
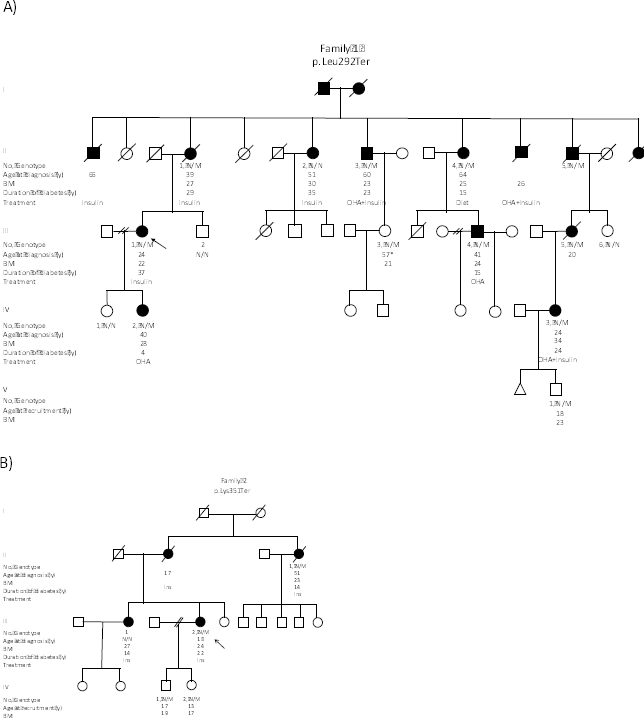
Extended pedigree of non–Finnish European patients identified in the discovery cohort. Genotype is shown underneath each symbol; M and N denote mutant and wild–type alleles, respectively. Directly below the genotype is the age of diabetes onset in years, duration in years, BMI and treatment at study entry. Squares represent male family members, and circles represent female members. Black–filled symbols denote patients with diabetes. An arrow denotes the proband in the family. OHA, oral hypoglycaemic agents. *age at recruitment. One of the daughters of patient lll.l in family 2 had a history of gestational diabetes.

**Table 1.** Frequency of heterozygous *RFX6* protein–truncating variants in all study cohorts and control populations.

|  | MODY cohorts,<br>Frequency of *RFX6* PTV | Control population,<br>Frequency of *RFX6* PTV | Odds ratio<br>(95% CI) | *P* |
| --- | --- | --- | --- | --- |
| Non-Finnish<br>European | Discovery cohort<br>(n=2/36), 5.5% | ExAC– exomes<br>(n=15/33346), 0.045% | 131<br>(14-595) | $1 \times 10^{-4}$ |
| | Replication cohort<br>(n=4/348), 1.15% | gnomAD– genomes<br>(n=2/7508), 0.027% | 43<br>(6-483) | $5 \times 10^{-5}$ |
| Finnish<br>European | Replication cohort<br>(n=6/80), 7.5% | METSIM- exomes<br>(n=26/7040), 0.37% | 22<br>(7-56) | $1 \times 10^{-6}$ |
| Meta-analysis<br>Heterogeneity chi-squared = 2.73 (d.f. = 2), *p* = 0.256 | | | 34<br>(15-80) | $1 \times 10^{-16}$ |

### Heterozygous *RFX6* PTVs were enriched in a MODY discovery cohort compared to population controls from ExAC

We compared the frequency of *RFX6* PTVs in our discovery cohort to a large control population with whole–exome data from ExAC^12^. Neither of the *RFX6* variants from the discovery cohort were present in the 60,706 individuals in ExAC. There were 15 individuals with *RFX6* PTVs in the 33,346 ExAC non–Finnish European control population (**Supplementary Table 3**).The frequency of the *RFX6* PTVs in the MODY discovery cohort was significantly higher (after accounting for the multiple testing of 29 genes) than the ExAC non–Finnish European control population (5.5% vs 0.045%, 131, 95% CI 14–595, P=l×l0^‐4^)(Table 1).

### Heterozygous *RFX6* PTVs are associated with MODY in a non–Finnish European replication cohort

To replicate the findings of our discovery cohort, we then examined 348 non–Finnish European probands who were routinely referred for MODY genetic testing to the Molecular Genetics Laboratory, Exeter, UK and in whom the common causes of MODY were excluded using targeted–NGS assay **(Supplementary Table 1)**.The analysis of heterozygous PTVs identified four unrelated probands with two novel *RFX6* nonsense variants (p.Gln25Ter, p.Arg377Ter) (**Supplementary Fig. 1, Supplementary Table 3**).Similarly, to the discovery cohort, the MODY replication cohort was enriched for *RFX6* PTVs compared to the ExAC non–Finnish European control population (1.15% vs 0.045%, odds ratio, OR=26, 95% CI 6–82, P=3×l0^‐5^) **(Supplementary Table 4)**.This association was maintained when compared to an independent non–Finnish European control population with whole–genome sequence data from gnomAD (http://gnomad.broadinstitute.org) (Table 1, **Supplementary Table 4**).The frequency of *RFX6* PTVs in the gnomAD genome dataset (0.027%) is not statistically different to that in ExAC (0.045%, P=0.76).

### Finnish individuals had ~10–fold higher frequency of *RFX6* PTVs compared to non–Finnish Europeans

The ExAC database showed a relative abundance of *RFX6* PTVs in Finnish Europeans (15/3305, 0.45%) compared to non–Finnish Europeans (15/33,346, 0.045%) (**Supplementary Table 3**). All of the Finnish individuals in ExAC with *RFX6* PTVs had the same variant, p.His293Leufs. To further validate this finding in a larger Finnish control population, we analysed *RFX6* PTVs in 7040 control individuals from the METSIM study in Eastern Finland^16^. There were 26 individuals with *RFX6* PTVs in this cohort and all had the p.His293Leufs variant. The frequency of p.His293Leufs was not significantly different from the ExAC Finnish population frequency (0.37% vs 0.45%, P=0.63) (**Supplementary Table 3**). The METSIM study has contributed to the ExAC Finnish cohort, so to prevent duplication we used the data from the larger METSIM study for further analysis^12^.

### The *RFX6* p.His293Leufs variant is strongly enriched in Finnish MODY patients with unknown aetiology

To assess whether the p.His293Leufs variant is associated with MODY in Finnish patients, we genotyped the *RFX6* p.His293Leufs variant in 80 Finnish probands who were routinely referred for MODY genetic testing to Genome Center of Eastern Finland, University of Eastern Finland and did not have mutations in the most common MODY genes *(GCK, HNF1A, HNF4A* and *HNF1B)* (**Supplementary Table 1**).We identified six probands with the p.His293Leufs variant. The frequency of this variant was significantly higher in the Finnish MODY cohort compared to the METSIM controls (7.5% vs 0.37%, OR=22, 95% CI 7¬56, P=l×l0^‐6^) (Table 1).The meta–analysis of the three independent case–control analyses confirmed the strong association of *RFX6* PTVs with MODY in the study cohorts (OR=34, 95% CI 15–80, *P*=l×l0^‐16^) (Table 1).

### The enrichment of *RFX6* PTVs in unknown MODY individuals is not due to technical artefacts

To ensure that the association we observed is not due to differences in sequencing technologies or analysis pipelines between cases and controls, we performed a series of sensitivity analyses. This included comparisons to additional whole exome, whole genome and in–house control cohorts and an analysis that removed exon 1 which was the least well covered exon in ExAC. These sensitivity analyses (**Supplementary Table 4**) show that results are consistent for all these analyses.

### *RFX6* PTVs co–segregation with diabetes is consistent with reduced penetrance

To further assess the causality of *RFX6* PTVs, we conducted a co–segregation analysis in families with genetic data available on more than three affected individuals. We had only one family (family 1) with >3 affected individuals with genetic data (Fig. 1)^17^. The analysis showed that the *RFX6* variant p.Leu292Ter co–segregated in 9 out of 10 individuals with diabetes (LOD score = 0.65, P=0.04). One individual without the *RFX6* variant had diabetes which is likely to be a phenocopy of type 2 diabetes considering the large pedigree, age of diagnosis and obesity (51 years, BMI 30 kg/m^2^). There were two family members with an *RFX6* variant but with normal HbA1c level at the time of study (18 and 57 years) suggesting that *RFX6* PTVs may have reduced penetrance.

### *RFX6* PTVs showed reduced penetrance compared to *HNF1A* and *HNF4A* MODY

To assess the penetrance of *RFX6* PTVs for diabetes compared to common causes of MODY, we combined data for all six non–Finnish European proband families. There were 18 *RFX6*heterozygotes of whom five had not developed diabetes at study entry. 27% (95% CI 11–58) developed diabetes by the age of 25 years and 78% (95% CI 55–95) by 51 years (Fig. 2). Two out of six probands did not have affected parents at study entry (**Supplementary Fig. 1**). The penetrance of diabetes for *RFX6* heterozygotes was substantially lower compared to pathogenic variants of *HNF1A* (70%, 95% CI 67–72 by the age of 25 years and 97%, 95% CI 96–98 by 50 years) and moderately lower than pathogenic variants of *HNF4A* (55%, 95% CI 50–60 by the age of 25 years and 91%, 95% CI 88–94 by 50 years) (Fig. 2). Similar to non– Finnish European proband families, the Finnish *RFX6* p.His293Leufs variant also showed reduced penetrance in Finnish families (**Supplementary Fig. 2**). In two previously reported families of neonatal diabetes children with homozygous p.Argl81Gln *RFX6*^15,18^ or *RFX6*p.His293Leufs^21^, the genetic information available on *RFX6* heterozygous family members was also suggestive of reduced penetrance of diabetes (**Supplementary Fig. 2, Supplementary Fig. 3**).

**Fig. 2.**
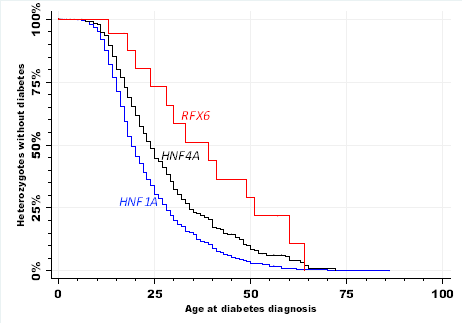
Penetrance of diabetes in people with a heterozygous RFX6 protein–truncating variant (n=18), pathogenic HNF1A variant (n=1265) or HNF4A variant (n=427).

### *RFX6* PTVs are not associated with type 2 diabetes

The reduced penetrance and later age of onset of diabetes with *RFX6* PTVs raised the possibility that these variants may be associated with type 2 diabetes. To assess this, we used freely available data from the Type 2 diabetes Knowledge Portal which contains whole–exome data on type 2 diabetes patients^19^. Burden testing of *RFX6* PTVs for exome sequencing data from 8373 type 2 diabetes cases and 8466 controls showed no significant association with type 2 diabetes (0.14% vs 0.083%, OR=1.79, 95% CI 0.7–4.57, *P=*0.22)^19^ **(Supplementary Table 3)**.

### Phenotype of RFX6–MODY

We assessed the diabetes phenotype in 27 *RFX6* heterozygote individuals with diabetes. The clinical features are shown in Table 2.The median age at diagnosis of diabetes was 32 years (IQR 24–46, range 13–64 years) and median BMI of 25.1 kg/m^2^ (IQR 23–28). After a median 10 years (IQR 5–22) of diabetes 69% of patients were treated with insulin but there was significant endogenous insulin present in 24/25 patients at recruitment. There was no history of sulphonylurea sensitivity and they did not have islet autoantibodies (GADA/IA2–Ab). All patients had isolated diabetes and there were no reports of the other features of homozygous *RFX6* mutations, such as duodenal or gall bladder atresia.

**Table 2.**
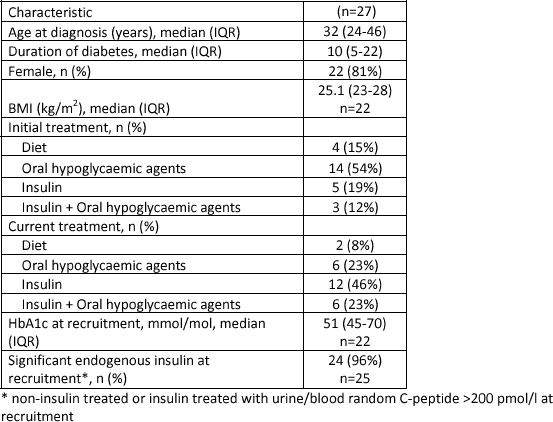
Clinical characteristics of patients with *RFX6-MODY*.

| Characteristic | (n=27) |
| --- | --- |
| Age at diagnosis (years), median (IQR) | 32 (24-46) |
| Duration of diabetes, median (IQR) | 10 (5-22) |
| Female, n (%) | 22 (81%) |
| BMI (kg/m<sup>2</sup>), median (IQR) | 25.1 (23-28)<br>n=22 |
| Initial treatment, n (%) |  |
| Diet | 4 (15%) |
| Oral hypoglycaemic agents | 14 (54%) |
| Insulin | 5 (19%) |
| Insulin + Oral hypoglycaemic agents | 3 (12%) |
| Current treatment, n (%) |  |
| Diet | 2 (8%) |
| Oral hypoglycaemic agents | 6 (23%) |
| Insulin | 12 (46%) |
| Insulin + Oral hypoglycaemic agents | 6 (23%) |
| HbA1c at recruitment, mmol/mol, median (IQR) | 51 (45-70)<br>n=22 |
| Significant endogenous insulin at recruitment*, n (%) | 24 (96%)<br>n=25 |

\* non-insulin treated or insulin treated with urine/blood random C-peptide >200 pmol/l at recruitment

### *RFX6* haploinsufficiency is associated with reduced gastric inhibitory polypeptide (GIP) secretion

*RFX6* is a transcription factor and has been shown to increase expression and secretion of GIP in mouse enteroendocrine K–cells^20^. We therefore measured the incretin hormone GIP in 17 *RFX6* heterozygotes (8 with diabetes) and compared to 26 controls (2 with diabetes). The fasting GIP was markedly lower in *RFX6* heterozygotes compared to controls (16 [10–24] vs 49 [28–65] pg/ml, *P=* 1.2×l0^5^). Fasting glucagon–like peptide–1 (GLP– 1) levels were not different in both groups (23 [12.5–32] vs 24 [14–32] pg/ml, *P=*0.98). To remove potential confounding factors we compared the OGTT data for the 11 Finnish *RFX6*p.His293Leufs heterozygotes without diabetes to five matched (age, sex, and BMI) controls for each heterozygote from the PPP–Botnia Study (Fig. 3, **Supplementary Table 6**). This confirmed that both fasting and 120 minute stimulated GIP was reduced (18.3 vs 48.9 pg/ml, P=8×l0^‐3^, 167 vs 241 pg/ml, *P=*0.029 respectively). In addition, the non–diabetic heterozygotes had higher fasting glucose (5.5 vs 5.1 mmol/l, *P=*0.02) with a similar fasting insulin level suggesting a beta–cell defect (**Supplementary Table 6**).

**Fig.3.**
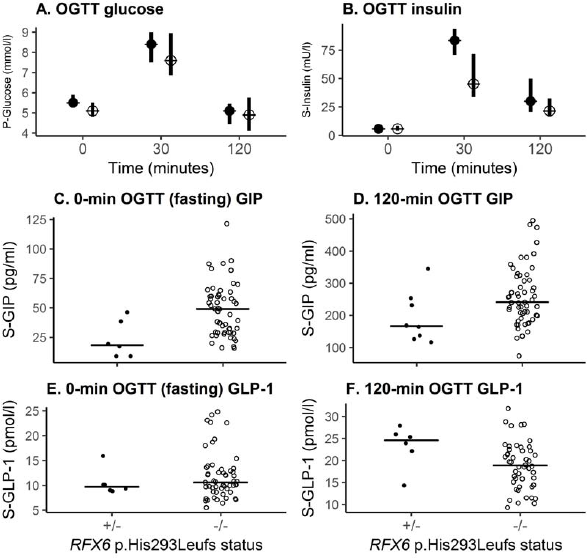
Phenotypic characteristics of the Finnish *RFX6* p.His293Leufs heterozygotes without diabetes. Filled symbol denotes *RFX6* heterozygotes (n=ll) and open symbol denotes population controls from the PPP–Botnia Study (1:5 matched for age, sex and BMI; N=55). Data are shown as medians (A–F) and interquartile range (A,B). *P* values <0.05 for (A) Glucose 0 min, *P*=0.02; (B) Insulin 30 min, *P*=0.01; (C) GIP 0 min, *P*=8xl0"^3^,120 min, *P*=0.02; (D) GLP–1120 min, *P*=0.047. *P* denotes plasma and S denotes serum. +/‐, Heterozygous for *RFX6* variant, –/–, without *RFX6* variant.

## Discussion

### Heterozygous *RFX6* protein–truncating variants are associated with MODY

We identified *RFX6* PTVs predicted to be pathogenic^11,21^ in unrelated MODY patients in whom the known causes of monogenic diabetes had been excluded. These variants were enriched in Finnish and non–Finnish European MODY probands but were rare in the control cohorts and in patients with type 2 diabetes. We observed co–segregation within pedigrees albeit with reduced penetrance. Finally, these variants are likely to have a functional effect due to nonsense mediated decay causing haploinsufficiency^12,22^. This is further supported by the studies that showed that the homozygous *RFX6* PTVs cause neonatal diabetes ^15,23,24^. Among unknown MODY cases, *RFX6* PTVs were responsible for 7.5% Finnish cases compared to only ~1% of non–Finnish European cases.

### Large–scale control cohorts such as ExAC in combination with next–generation sequencing of well–characterised cases is a useful strategy for identifying new causes of monogenic disease

The large ExAC database provides sufficient power for reliable burden testing of rare variants in monogenic disease^12^ without the need for large pedigrees or linkage analysis. *RFX6* PTVs were highly enriched in both discovery and replication cohorts compared to control cohorts, supporting their pathogenicity. The ExAC database has been very useful in identifying benign variants because of an unusually high frequency in the population compared to frequency of the disease in question ^15,23,24^. However, caution is required for reduced penetrance variants as their frequency can be higher than estimated disease frequency in the general population. *RFX6* PTVs are an example where the reduced penetrance explains the higher frequency in control cohorts compared to the estimated frequency of MODY (0.01%) in the general population^26^. In addition, our study highlights the importance of population specific control and disease cohorts. The frequency of *RFX6* PTVs is ~10–fold higher in the Finnish population compared to non–Finnish populations due to the well–documented bottleneck in population genetics^27,28^. This observation is not restricted to *RFX6* alone. It has been shown that the Finnish population has an overall higher burden of genome–wide PTVs (0.5–5%) in many genes compared to non–Finnish Europeans, and some of these have been associated with disease^27^–^29^.

### *RFX6* PTVs are associated with reduced penetrance MODY

This reduced penetrance explains the lack of complete co–segregation in the *RFX6* pedigrees. It also clarifies why diabetes was only reported in the parents or grandparents (obligate heterozygous for functional *RFX6* variants) in 7 out of the 12 published recessive *RFX6* neonatal diabetes pedigrees^15,18,24,30–36^. The lack of enrichment of *RFX6* PTVs in type 2 diabetes patients compared to controls further supports their association with reduced penetrance MODY rather than type 2 diabetes. Further studies are needed to understand the mechanism of reduced penetrance of diabetes in *RFX6* heterozygotes. It could be due to a combination of factors, such as expression patterns of normal alleles, epigenetic modifications and rare or common genetic variant modifiers^37^.

### There are differences as well as similarities between RFX6–MODY and *HNF1A/HNF4A–* MODY

In contrast to *HNF1A* and *HNF4A–MODY* patients, *RFX6–*MODY patients do not show enhanced sensitivity to sulphonylureas^38^. All patients with *HNF1A/HNF4A–MODY* have significant endogenous insulin 3–5 years post diagnosis^39^. *RFX6–*MODY patients showed a similar pattern, except for one patient who did not have detectable endogenous insulin. Similar to *HNF1A/HNF4A–MODY^6,38^, RFX6–*MODY patients lack islet autoantibodies and have isolated diabetes. This suggests that persistent C–peptide, lack of islet autoantibodies and parental history of diabetes which are currently used to distinguish common forms of MODY from type 1 diabetes, can also be used to identify *RFX6–*MODY. However, a similar strategy will not help to distinguish late onset *RFX6–*MODY from type 2 diabetes.

### Our study supports the role of RFX6 in the adult human pancreas

RFX6 is from a family of transcription factors that contains winged–helix DNA binding domains^15^. RFX6 is expressed almost exclusively in pancreatic islets, small intestine and colon^15^. It acts downstream of NGN3, regulates islet cell differentiation and the development of the endocrine pancreas^15^. The homozygous *RFX6* missense and PTVs cause syndromic neonatal diabetes (gall bladder aplasia, gut atresia and diabetes)^15^. *RFX6* whole–body null mice show phenotypes consistent with human disease and die soon after birth, but the heterozygous whole–body *RFX6* mouse has not been reported to develop diabetes^15^. This is not surprising considering the lack of phenotype in heterozygous null mice of *HNF1A* and *HNF1B*^40^. Interestingly, the defect in glucose induced insulin secretion was present in models that are more akin to haploinsufficiency of *RFX6* in adult humans^33,41^. 80% depletion of RFX6 protein in the adult mouse pancreas *in vivo* as well as in human beta cells *in vitro* showed that this defect was due to reduced expression of ABCC8, GCK and Ca^2+^channels in beta cells and disruption of Ca^2+^ mediated insulin secretion^33,41^. These data support the role of RFX6 in the physiology of adult beta cells. This along with evidence of impaired of beta–cell function (requirement of insulin to maintain euglycemia, one patient with C–peptide <200 pmol/l, lower C–peptide in heterozygotes with diabetes compared to without diabetes **[Supplementary Fig. 4]**), suggest that insulin deficiency is the cause of diabetes in these patients.

### A role for RFX6 in the secretion of the incretin hormone GIP

Incretins are gut hormones released in response to meals that potentiate glucose–stimulated insulin secretion. GIP is secreted from enteroendocrine K–cells in the duodenum and upper jejunum, and mediates the bulk of the incretin effect in healthy individuals^42^. The secretion of GIP and GLP–1 is preserved in type 2 diabetic patients^43,44^ and in patients with other forms of diabetes, including type 1 diabetes^45^ and *HANF1A–MODY*^46^. The present identification of GIP deficiency in *RFX6* PTV heterozygotes is in keeping with the murine data showing that GIP expression and secretion is regulated by Rfx6^20^, and, importantly, identifies the first human form of diabetes associated with decreased GIP secretion.

In conclusion, heterozygous *RFX6* PTVs are associated with reduced penetrance MODY and GIP deficiency.

## Methods

### Study populations

*Discovery MODY cohort:* The discovery cohort comprises of 38 European probands with strong MODY–like phenotype who did not have mutations in the three most common MODY genes *(GCK, HNF1A* and *HNF4A*) (**Supplementary Table 1**). They were diagnosed <25 years of age, non–obese, had ≥ 3 generation history of diabetes, non–insulin treated or insulin treated with C–peptide > 200 pmol/L (if available), and lacked islet autoantibodies.

*Non–Finnish European replication MODY cohort:* The replication cohort was derived from 469 non–Finnish European routine MODY diagnostic referrals to the Molecular Genetic Laboratory, Exeter, UK. A monogenic aetiology in a known monogenic diabetes gene was identified in 121 patients and the remaining 348 patients with unknown aetiology comprised the replication cohort **(Supplementary Table 1)**.

*Finnish–European replication MODY cohort:* This cohort consisted of 80 patients who were routinely referred for MODY diagnostic testing to the Genome Center of Eastern Finland, University of eastern Finland in whom no mutation was found in the common MODY genes *(GCK, HNF1A, HNF1B* and *HNF4A)* when assessed by Sanger sequencing **(Supplementary Table 1)**. These 80 patients comprise 78% of the total MODY X Finnish cohort.

*Finnish–European control cohort:* Individuals of this cohort were part of the METSIM study (n=7040). They were all males aged 45 to 70 years, randomly selected from the population register of the Kuopio town, Eastern Finland and have been described previously^16^.

*Cohort of people with pathogenic HNF1A and HNF4A variants:* This cohort included probands and their family members referred to the Molecular Genetics Laboratory, Exeter, UK for MODY genetic testing and were identified to have a pathogenic *HNF1A* (n=1265) or *HNF4A* (n=427) variant.

*Phenotypic characterization of RFX6 heterozygotes:* In total, we had 47 *RFX6* heterozygotes of whom 27 had diabetes. 29/47 were part of the discovery and replication cohorts. 18/47 were identified separately or had been previously reported^18,23^ **(Supplementary Fig. 2–Family 3–5, Supplementary Fig. 3)**. The clinical features of *RFX6–MODY* were based on 27 individuals with diabetes. 13/27 were part of the discovery and non–Finnish replication cohort **(Fig. 1, Supplementary Fig. 1–Family 1–6)**. 9/27 individuals were from the Finnish replication cohort (5/9 individuals were from **Supplementary Fig. 2–Family 1 and 2**, Pedigrees were not available for 4/9 individuals). In addition to this, we included 5 diabetic individuals that were identified separately. This comprised a single Finnish individual **(Supplementary Fig. 2–Family 3)** and 4 individuals from a previously reported family from Belgium **(Supplementary Fig. 3)** ^18^.

*Incretin analysis:* We completed physiological studies on 25/47 *RFX6* heterozygotes in whom 10 had diabetes. 7/25 had fasting blood sample analysis and 18/25 had 75g oral glucose tolerance test (OGTT). We used 27 (12 family and 15 unrelated) controls that included 2 individuals with diabetes for an initial comparison of fasting GIP/GLP–1. *RFX6* heterozygotes had similar age (38 [ 34–58] vs 40 [ 30–61] years, P=0.70), sex (female 64% vs 56% *P=*0.58) and BMI (24 [23–29] vs 27[24–30] kg/m^2^, P=0.27) distribution as the 27 controls. Fasting GIP/GLP–1 levels was available on 17/25 *RFX6* heterozygotes and 26/27 controls. Out of 25 individuals who were assessed for GIP/GLP–1 analysis, 16 Finnish individuals (5 with diabetes, 11 without diabetes) participated in the FINN MODY Study(http://www.botnia–studv.org/finnmodv, recruiting patients with MODY–like diabetes and their relatives in Finland). It is linked to the population–based PPP–Botnia study and the participants had been subjected to standardised OGTT, sample collection and detailed biochemical analysis following the study protocol of the PPP–Botnia study. Therefore, to remove any potential confounding factors, we compared the phenotypic characteristics of these 11 *RFX6*p.His293Leufs heterozygotes without diabetes to five controls for each heterozygote from the PPP–Botnia study matched for age, sex and BMI. None of the controls had the *RFX6*p.His293Leufs variant.

*PPP–Botnia Study:* The Prevalence, Prediction and Prevention of diabetes (PPP)–Botnia Study is a population–based study in Western Finland aiming at obtaining accurate estimates of prevalence and risk factors for T2D, impaired glucose tolerance, impaired fasting glucose and the metabolic syndrome in the adult population (Isomaa). Altogether 5208 individuals randomly recruited from the national Finnish Population Registry participated in the baseline study in 2004–2008 (representing 6–7% of the population), and 3870 (77%) individuals participated in the follow–up study in 2011–2014. The participants with fasting plasma glucose < 10 mmol/l participated in an 75g OGTT with venous samples taken at 0, 30,120 min for plasma glucose and serum insulin; at 0 and 120 min for serum C–peptide, GIP and GLP–1. The participants gave their written informed consent and the study protocol was approved by the Ethics Committee of Helsinki University Hospital, Finland.

Plasma glucose was analyzed using the Hemocue Glucose System (HemoCue AB, Ängelholm, Sweden). Serum insulin was measured by an AutoDelfia fluoroimmunometric assay (PerkinElmer, Waltham, Massachusetts, US) and serum C–peptide by Cobas e411 electrochemiluminometric immunoanalysis (Roche, Mannheim, Germany). Serum GIP was analyzed using Millipore's Human GIP Total ELISA (Merck, Darmstadt, Germany; catalogue # EZHGIP–54K), which has 100% cross–reactivity to both human GIP (1–42) and GIP (3–42). Serum total plasma GLP–1 concentrations, which detects both intact GLP–1 and GLP–1 (9–36 amide), were determined using Millipore's radioimmunoassay (Merck, Darmstadt, Germany; catalogue #GLP1T–36HK. Serum total cholesterol, HDL and triglyceride concentrations were measured first on a Cobas Mira analyzer (Hoffman LaRoche, Basel, Switzerland) and LDL cholesterol concentrations were calculated using the Friedewald formula, and since January 2006 with an enzymatic method (Konelab 60i analyser; Thermo Electron Oy, Vantaa, Finland).

*The RFX6>* p.His293Leufs variant was genotyped in 5187 individuals by the Kompetitive Allele Specific PCR genotyping system (KASPTM) on Demand (KOD) assay according to the manufacturer's testing conditions including six positive control samples identified by direct sequencing (LGC Hoddesdon, Herts, UK). Two out of 5180 participants had *RFX6*p.His293Leufs (the genotyping failed in 7), which was confirmed by direct sequencing.

### DNA analysis

#### Targeted–next generation sequencing

The analysis of all known monogenic diabetes genes in European cohorts was conducted using targeted–NGS as described previously^13^. The panel included 29 genes in which variants are known to cause monogenic neonatal diabetes, MODY, mitochondrial diabetes, lipodystrophy or other forms of syndromic diabetes^13^ **(Supplementary Table 2)**. The *RFX6*PTVs identified by targeted–NGS were confirmed using Sanger sequencing. The essential splice site, nonsense and frameshift variants excluding the last exon were considered PTVs in this study^12,22^. The targeted–NGS assay covered 100% bases of the *RFX6* coding region >10x read depth for all the samples.

#### Sanger sequencing

Genomic DNA was extracted from whole blood using standard procedures and the coding region and intron/exon boundaries of the *RFX6* gene were amplified by PCR. Amplicons were sequenced using the Big Dye Terminator Cycler Sequencing Kit v3.1 (Applied Biosystems, Warrington, UK) according to manufacturer’s instructions and reactions were analysed on an ABI 3730 Capillary sequencer (Applied Biosystems, Warrington, UK). Sequences were compared with the reference sequences (NM_173560.3) using Mutation Surveyor v3.24 software (So Genetics, State College, PA, USA).

The Finnish–European MODY cohort was analysed for p.His293Leufs variant using Sanger sequencing as described above. Family co–segregation analysis was performed in available family members using a Sanger sequencing assay for the specific *RFX6* variant identified in that family. DNA analysis of the METSIM study has been described before^23^.

### Statistical analysis

Fisher’s exact test was used to compare the frequency of *RFX6* PTVs. The threshold p value for association was l×l0^‐3^ as there were 29 genes on the panel (0.05/29). The penetrance of diabetes was assessed using survival time analysis method. The statistical analysis was conducted using Stata 14 (StataCorp, Texas, USA). The comparison of *RFX6* heterozygotes to PPP–Botnia controls were conducted using R (3.3.2) with packages for the data manipulation (dplyr) and visualization (ggplot2). Continuous variables were compared with Mann– Whitney U test and categorical variables with chi–squared test. Single point non–parametric linkage analyses were performed using MERLIN 1.1.2^47^. The Z score was converted into a LOD score by use of the Kong and Cox exponential model implemented in MERLIN^47,48^.

### Ethics

The informed consent was obstained from all subjects. The study is approved by the North Wales Research Ethics Committee, UK. The FINNMODY/PPP–Botnia study is approved by the Research Ethics committee for Medicine of the Helsinki University Hospital.

#### Duality of Interest

No potential conflicts of interest relevant to this article were reported.

#### Author Contributions

K.A.P., J.K., M.C. and M.N.W. researched data and performed statistical analyses. K.A.P., J.K., M.C. and T.T. wrote the first draft of the manuscript, which was modified by all authors. All authors contributed to the discussion and reviewed or edited the manuscript. M.N.W., M.C. and T.T. are the guarantors of this work and, as such, had full access to all the data in the study and take responsibility for the integrity of the data and the accuracy of the data analysis.

#### Funding

K.A.P. has a postdoctoral fellowship funded by the Wellcome Trust (110082/Z/15/Z). S.E.F. has a Sir Henry Dale Fellow–ship jointly funded by the Wellcome Trust and the Royal Society (105636/Z/14/Z). S.E. and A.T.H. are Wellcome Trust Senior Investigators (WT098395/Z/12/Z), and A.T.H. is also supported by a NIHR Senior Investigator award. M.N.W. is supported by the Wellcome Trust Institutional Strategic Support Fund (WT097835MF) and the Medical Research Council (MR/M005070/1). M.S. is supported by the NIHR Exeter Clinical Research Facility. Additional support came from the University of Exeter and the NIHR Exeter Clinical Research Facility. M.C. is supported by the European Union’s Horizon 2020 research and innovation programme, project T2DSystems, under grant agreement No 667191, and the Fonds National de la Recherche Scientifique (FNRS) and Actions de Recherche Concertées de la Communauté Française (ARC), Belgium. The FINNMODY and PPP–Botnia Studies have been financially supported by the Sigrid Juselius Foundation, the Folkhalsan Research Foundation, Helsinki University Central Hospital Research Foundation, Finnish Diabetes Research Foundation, Finnish Medical Society, Foundation for Pediatric Research, Ahokas Foundation, Ollqvist Foundation, Nordic Center of Excellence in Disease Genetics, EU–EXGENESIS), Signe and Ane Gyllenberg Foundation, Swedish Cultural Foundation in Finland, Finnish Diabetes Research Foundation, Foundation for Life and Health in Finland, Paavo Nurmi Foundation, Perklén Foundation, Närpes Health Care Foundation, and Diabetes Wellness Foundation. The study has also been supported by the Municipal Heath Care Center and Hospital in Jakobstad, Health Care Centers in Vasa,Narpes and Korsholm. The views expressed are those of the authors and not necessarily those of the Wellcome Trust, the National Health Service, the NIHR, or the Department of Health.

